# One of these morphs is not like the others: orange morphs exhibit different escape behavior than other morphs in a color polymorphic lizard

**DOI:** 10.1101/2021.12.14.472706

**Authors:** Kinsey M. Brock, Indiana E. Madden

## Abstract

Variation in color morph behavior is an important factor in the maintenance of color polymorphism. Alternative anti-predator behaviors are often associated with morphological traits such as coloration, possibly because predator-mediated viability selection favors certain combinations of anti-predator behavior and color. The Aegean wall lizard, *Podarcis erhardii*, is color polymorphic and populations can have up to three monochromatic morphs: orange, yellow, and white. We investigated whether escape behaviors differ among coexisting color morphs, and if morph behaviors are repeatable across different populations with the same predator species. Specifically, we assessed color morph flight initiation distance (FID), distance to the nearest refuge (DNR), and distance to chosen refuge (DR) in two populations of Aegean wall lizards from Naxos island. We also analyzed the type of refugia color morphs selected and their re-emergence behavior following a standardized intrusion event. We found that orange morphs have different escape behaviors from white and yellow morphs, and these differences are consistent in both of the populations we sampled. Orange morphs have shorter FIDs, DNRs, and DRs, select different refuge types, and re-emerge less often after an intruder event compared to white and yellow morphs. Observed differences in color morph escape behaviors support the idea that morphs have evolved alternative behavioral strategies that may play a role in population-level morph maintenance and loss.

## INTRODUCTION

Understanding how phenotypic variation is generated and maintained is a central goal in evolutionary biology. Color polymorphic species are good systems for studying mechanisms that regulate phenotypic variation because color morphs represent distinct, genetically-determined variation maintained by selection (Gray & McKinnon 2007; Svensson 2017). Color morphs usually, if not always, differ in additional traits (McKinnon & Pierotti 2010; Stuart-Fox et al. 2021) such as morphology (Brock et al. 2020), physiology (Border et al. 2019), diet (Lattanzio & Miles 2016), and behavior (Vercken & Clobert 2008). These alternative morph phenotypes are likely the result of correlational selection that produces genetic correlations between phenotypic traits under selection, such as color and behavior (Brodie 1989; Brodie 1992).

Predation can be a major selective force in the evolution of color and behavior (Brodie 1992; Lima & Dill 1990; Hurtado-Gonzales et al. 2010; Pérez i de Lanuza & Font 2015), and could play a role in maintenance of color polymorphism. Because color is a visual trait, certain color morphs could be more easily detected by predators and may use different tactics to avoid predation (Forsman & Appleqvist 1998). In insects (Forsman & Appleqvist 1998), fish (Maan et al. 2008) and reptiles (Forsman 1995), intra-specific color morphs suffer different rates of predation in different environments due to differential visual detection by predators. While predators refine their hunting techniques to find and obtain prey, prey evolve adaptive strategies to avoid being detected and captured (Dawkins & Krebs 1979). For color morphs, avoiding detection by predators may require different strategies given differences in conspicuousness against different backgrounds and surroundings (Marshall et al. 2016). Prey that differ in color or pattern may evolve different anti-predator behaviors that involve occupying distinct microhabitat to enhance crypsis and avoid detection (Reid 1987; Sandoval 1994), different levels of boldness (Yewers et al. 2016), and different escape behaviors (Marcellini & Jenssen 1991; Labra et al. 2007).

All prey species must balance the risk of predation with activities that allow them to meet their basic needs, and color morphs may evolve different strategies for negotiating safety based on their risk of being visually detected by predators. The flight initiation distance (FID) is the distance between a prey animal and a perceived threat when the prey animal begins its escape (Ydenberg & Dill 1986; Runyan & Blumstein 2004). Generally, FIDs have been used as a metric of risk assessment and habituation (Blumstein 2003), where longer FIDs are interpreted as risk aversion, and shorter FIDs are associated with boldness or habituation (Blumstein 2006). FIDs have also been useful for quantifying variation in escape behavior and capture the benefits and costs to fleeing that prey animals must negotiate (Lima & Dill 1990). If prey flee at greater distances upon perceiving a predation threat, they may lose out on beneficial feeding, mating, social, or thermoregulatory opportunities that can increase prey fitness (Lima & Dill 1990). On the other hand, prey must weigh the benefits of staying against the costs of staying too long when faced with a predation threat. Costs of staying too long can include an energy-extensive chase, injury, or death by predation. Ultimately, escape is predicted to begin when a predator approaches to a distance where the cost of staying and the cost of fleeing are equal (Ydenberg & Dill 1986; Cooper & Frederick 2007). Since prey usually flee to a refuge (a space a predator can not access), deciding when to flee may be influenced by the distance a prey is from the nearest refuge (DNR). Prey must also quickly select what type of refuge to take and decide how long to wait to safely re-emerge at the expense of other activities. If predation risk is unequal among morphs that differ in color, we expect morphs to display consistent, distinct escape behaviors when confronted with a simulated predation event.

The present study focuses on color morph escape behavior in the color polymorphic Aegean wall lizard (*Podarcis erhardii*). This species has three co-occurring monochromatic color morphs that are either orange, yellow, or white. All sexes are color polymorphic in this species and can be any of the three colors (Brock et al. 2020). Male color morphs differ in head and body size and female color morphs tend to be more similar in size (Brock et al. 2020). Male color morphs exhibit differences in several social behaviors like aggression, boldness, chemical signaling, and visual signaling frequency when in contest with each other over limited resources (Brock et al. 2021a in review), suggesting that color morphs in this species may have different behavioral strategies. We focused our study on color morph risk assessment and escape behavior, including flight initiation distance (FID), distance to the nearest refuge (DNR), refuge choice (distance to chosen refuge and refuge type), and re-emergence behavior. In this study, we address the following questions:

1. Do color morphs differ in their escape behavior (flight initiation distance, distance to nearest refuge, and distance to chosen refuge) and are these morph differences in behavior consistent across two populations with the same predator guilds?
2. Do color morphs differ in the type of refuge they select when fleeing from an approaching threat?
3. After taking refuge, do color morphs differ in the amount of time they wait to re-emerge from hiding?

To answer these questions, we measured the escape behavior of color morphs observed on dry stone walls from two populations on Naxos. We hypothesized that color morphs within a site would exhibit significantly different escape behaviors (FID, DNR, DR, refuge selection, and re-emergence), and that these differences would be consistent among populations with the same predator guilds. Specifically, we expected white morphs to have shorter FIDs and longer DNRs and DRs based on *ex situ* behavioral experiments that revealed white morphs behave more aggressively and boldly than other morphs (Brock et al. 2021a in review). We also expected orange morphs to have longer FIDs and shorter DNRs because it was determined that orange morphs in a closely related polymorphic *Podarcis* species were more conspicuous to conspecifics and predators than yellow or white morphs (Pérez i de Lanuza & Font 2015). We hypothesized that color morphs would differ in the types of refuge they select when fleeing from similar starting positions on dry stone walls. Here, we expected that orange morphs would seek refuge in vegetation more often than other color morphs since they are most often found in microhabitat that is shaded by leafy vegetation (Brock, unpublished data). Finally, we hypothesized that color morphs would differ in their re-emergence behavior.

Specifically, we expected white and yellow morphs to re-emerge from hiding more often than orange morphs, and take less time to do so, given their tendency to be more active, aggressive, and bold than orange morphs (Brock et al. 2021a in review).

## METHODS

### Study sites

We conducted our study in June 2021, in the middle of the breeding season when lizards are most active (Valakos et al. 2008). We measured the escape behavior of color morphs in two populations from the island Naxos: Moni and Damarionas. Moni and Damarionas are separated by the Halki valley and a distance of 7 km apart. Both sites have all three color morphs and are similar environmentally and ecologically. Moni and Damarionas are small agricultural villages with olive groves, other fruit bearing trees, and small fields primarily used for goat grazing. Both sites contain many dry stone walls that wall lizards occupy to shelter from predators, safely thermoregulate, and forage for invertebrates. The natural vegetation is characteristic of Mediterranean scrub habitat with sclerophyllous evergreen maquis, and *phrygana*, a diverse species community of summer-deciduous shrubs (Fielding and Turland 2008). The predator guild is the same at both sites where snakes (*Elaphe quatuorlineata, Viper ammodytes, Natrix natrix*, and *Eryx jaculus*), small mammals (*Rattus rattus, Martes foina*), introduced feral house cats, (*Felis catus*), and birds (*Falco tinnunculus* and *Buteo buteo*) prey upon *P. erhardii* (Brock et al. 2015).

### Field sampling of escape behavior

Field sampling took place on warm, sunny days with low wind speeds when lizards were fully active. We recorded flight initiation distance (FID) using a method developed by Blumstein (2003) and utilized in many previous studies on various lizard species including *P. erhardii* (Li et al. 2014; Brock et al. 2015). To eliminate experimenter bias, the same person performed all FID approaches in the field while wearing the same clothes. We limited our sampling to lizards initially spotted on dry stone walls so that comparisons of morph escape behavior and refuge choice come from the same starting conditions. The experimenter walked through each study site at a practiced pace (80 m/min [Pérez-Cembranos et al. 2013]) until they spotted a lizard.

Upon sight, the experimenter would confirm a lizard’s color morph identity and sex with binoculars. Color morphs in this species can be reliably distinguished by eye (Brock et al. 2020), and females and males are easily distinguishable by their head and tail shapes. Once the color morph and sex of the individual was recorded, the experimenter walked directly toward the focal lizard from a starting distance of approximately 500 cm away. Approaches were performed at a practiced pace (0.5 m/s) while maintaining eye contact and the experimenter stopped once the focal lizard moved away (i.e.: initiated an escape response). The experimenter then recorded the distance between their stopping point and the point of lizard flight initiation (FID). The experimenter also recorded the distance from the point of flight initiation and the nearest available refuge (distance to nearest refuge, DNR) and the distance from the point of flight initiation and where the lizard chose to seek refuge (distance to refuge, DR). We also recorded the type of refuge the lizard selected, which ultimately fell into one of two categories: either a crevice in the stone wall or vegetation. Finally, after each lizard entered a refuge, the experimenter waited 30 seconds and recorded whether or not the lizard re-emerged in that time frame and recorded the time in seconds.

To ensure we did not measure the same individual twice, we recorded our transect tracks with a Garmin GPS unit using the waypoint finder feature, so we did not revisit the same area while collecting FIDs. The home range size of this species is small (BeVier et al. 2021), and we did not record individuals of the same sex and morph very near to each other, so we think it is unlikely that we sampled the same individual twice.

### Ethical note

All procedures in this study were approved by the Greek Ministry for Environment and Energy (permit YΠEN/ΔΔΔ/43912/1357).

### Statistical analyses

Flight initiation distance can be influenced by the starting distance of the experimenter (Blumstein 2003). Therefore, we first tested for a statistical relationship between FID and starting distance to determine whether we needed to include starting distance as a covariate in our comparisons of FID. We did not find a significant linear relationship between FID and experimenter starting distance in our dataset (Pearson corr = 0.025, df = 418, p = 0.61), and proceeded to analyze morph differences in FID without starting distance as a covariate.

To test for color morph differences in mean flight initiation distance, distance to nearest refuge, and distance to refuge, we used a combination of Levene’s Tests, One-way ANOVAs, and Welch’s One-way ANOVAs. First, we assessed measurements of FID, DNR, and DR for normality and homogeneity of variance using Q-Qplots and Levene’s Tests. All data were normally distributed, but some groups had unequal variances. For groups that had equal variances, we used One-way ANOVAs to test for mean differences in escape behaviors. For groups that had unequal variances, we tested for differences in FID, DNR, and DR using Welch’s One-way tests. To determine which groups differed from each other, we followed up One-way ANOVAs and Welch’s One-way tests with post-hoc Tukey HSD and Games-Howell tests, respectively.

We tested if females and males of the same morph color from the same population had significantly different FIDs, DNRs, and DRs using t-tests. We determined if females and males had equal variances in FID, DNR, and DR using F tests and then used Two Sample t-tests for groups with equal variances, or a Welch’s Two sample t-test for groups with unequal variances to test for mean differences in escape behaviors.

To test for color morph associations with refuge selection (e.g. stone wall crevice or vegetation), we used chi-squared tests of independence.

We tested for color morph differences in re-emergence behavior using Kruskal-Wallis tests. To determine which groups differed significantly from each other, we used a Mann-Whitney U post hoc test. To prevent the inflation of type I error rates, or rejecting the null when the null is true, adjustments to the *p*-values were made using a Bonferroni correction, in which the p-values are multiplied by the number of comparisons.

We performed all statistical analyses in R (R Core Team 2021). Code and data for this project are available on Dryad.

## RESULTS

In total, we measured the escape behavior of 420 lizards. At Moni, we measured 50 of each monochromatic morph per sex, for a total of 300 lizards. At Damarionas, we measured 20 of each morph per sex for a total of 120 lizards. Flight initiation distance ranged from 5 to 206 cm (mean = 78.2 cm, ± 41.2 cm) at Moni, and from 30 to 187 cm (mean = 84.6 cm, ± 31.9 cm) at Damarionas. Distance to nearest refuge ranged from 0 to 50 cm at Moni (mean = 7.96 cm, ± 6.06 cm), and from 0 to 29 cm (mean = 7.59 cm, ± 6.47 cm) at Damarionas. Distance to chosen refuge ranged from 0 to 109 cm (mean = 15.37 cm, ± 15.69 cm) at Moni, and from 0 to 126 cm (mean = 16.11 cm, ± 21.6 cm) at Damarionas.

### Morph differences in escape behavior by site

Color morphs exhibited differences in escape behaviors, and these morph differences were consistent across both populations we sampled. Across both populations and all escape behaviors we measured, orange morphs always differed from the other two morphs (Figure 1).

**Figure 1.**
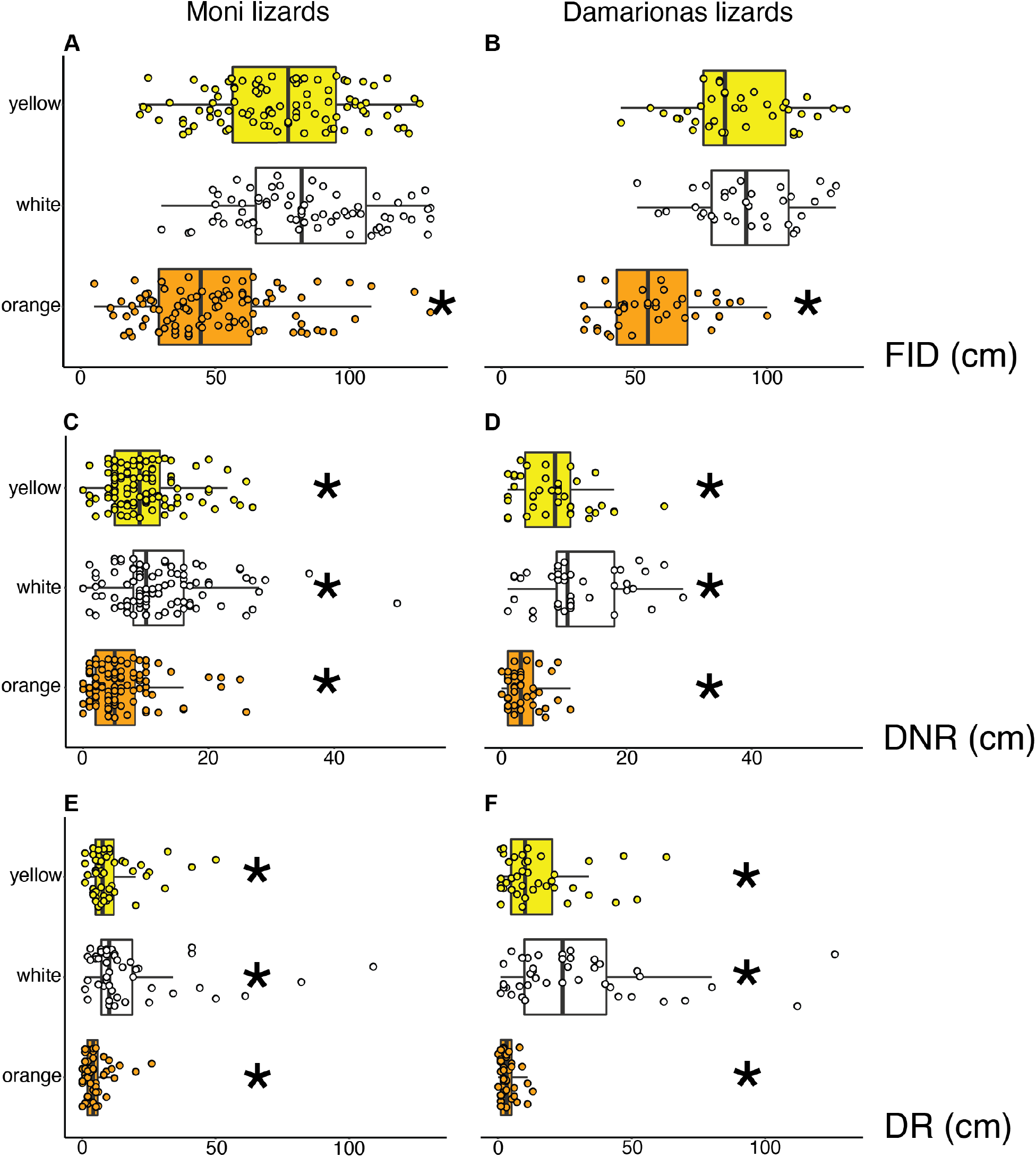
Color morph differences in flight initiation distance FID (A,B), distance to nearest refuge DNR (C,D), and distance to chosen refuge DR (E,F) at Moni and Damarionas. Asterisks indicate mean differences from all other color morphs (alpha < 0.05).

At both Moni and Damarionas, we found that orange morphs had significantly shorter FIDs than yellow and white morphs, which did not differ from each other (Table 1, Figure 1). We also found consistent differences across sites in the distance color morphs were observed from the nearest available refuge (DNR). At both Moni and Damarionas, we found all morphs differed from each other in their DNR (Figure 1). At both sites, orange morphs had the shortest DNR and white morphs had the longest DNR (Table 1). Finally, we found consistent color morph differences in the distance to their chosen refuge (Figure 1). Again at both sites, orange morphs had the shortest DR and white morphs had the longest DR (Table ANOVA).

**Table 1.** Test statistics for analyzing mean differences in morph escape behaviors. At both sites, orange morphs always differed from other morphs in their escape behavior.

| Site | Escape behavior Test | Post hoc comparisons | Difference in Means | *P* adj |
| --- | --- | --- | --- | --- |
| Moni | Morph FID<br>Welch's ANOVA<br>$F(2,190.9) = 75.72$ | orange-yellow<br>yellow-white<br>white-orange | -30<br>-6.8<br>58 | **< 0.001\***<br>=0.07<br>**< 0.001\*** |
| | Morph DNR<br>Welch's ANOVA<br>$F(2,193.3) = 23.75$ | orange-yellow<br>yellow-white<br>white-orange | -3.7<br>-2.5<br>6.2 | **< 0.001\***<br>**< 0.001\***<br>**<0.05\*** |
| | Morph DR<br>Welch's ANOVA<br>$F(2,164.4) = 36.07$ | orange-yellow<br>yellow-white<br>white-orange | -7.4<br>-8.2<br>15.6 | **< 0.001\***<br>**< 0.001\***<br>**< 0.001\*** |
| Damarionas | Morph FID<br>ANOVA<br>$F(2,117) = 37.63$ | orange-yellow<br>yellow-white<br>white-orange | -34.17<br>-8.6<br>45.77 | **< 0.001\***<br>= 0.28<br>**< 0.001\*** |
| | Morph DNR<br>Welch's ANOVA<br>$F(2,66.1) = 31.92$ | orange-yellow<br>yellow-white<br>white-orange | -4.8<br>-3.8<br>8.6 | **< 0.001\***<br>**< 0.001\***<br>**< 0.001\*** |
| | Morph DR<br>Welch's ANOVA<br>$F(2,54.74) = 25.84$ | orange-yellow<br>yellow-white<br>white-orange | -11.4<br>-14.7<br>26.1 | **< 0.001\***<br>**< 0.001\***<br>**< 0.001\*** |

### Sex differences in escape behavior by morph

At Moni, orange and yellow morphs exhibited sex differences in flight initiation distance, while morphs did not (Figure 2 A, B, C). Orange females had significantly longer FIDs than orange males on average (Welch’s t = 8.73, df = 82.61, p < 0.001), and yellow morphs followed the same pattern (t = 2.24, df = 98, p = 0.03). We observed no significant sex differences in FID for the white morph at Moni (t = 1.35, df = 98, p = 0.18). At Moni, we found that all morphs exhibited sex differences in distance to the nearest refuge (Figure 2 D, E, F). Orange males were spotted significantly further away from the nearest refuge than orange females (t = −4.05, df = 98, p < 0.001), and the same pattern was observed in white morphs (t = −4.01, df = 98, p < 0.001), and yellow morphs (t = −3.52, df = 98, p < 0.001). Finally, at Moni, all morphs exhibited sex differences in the distance to chosen refuge (Figure 2 G, H, I). Orange (t = −4.96, df = 98, p < 0.001), white (t = −2.88, df = 98, p = 0.005), and yellow (t = −3.21, df = 98, p = 0.002), females all chose significantly closer refuges than males of their same color morph.

**Figure 2.**
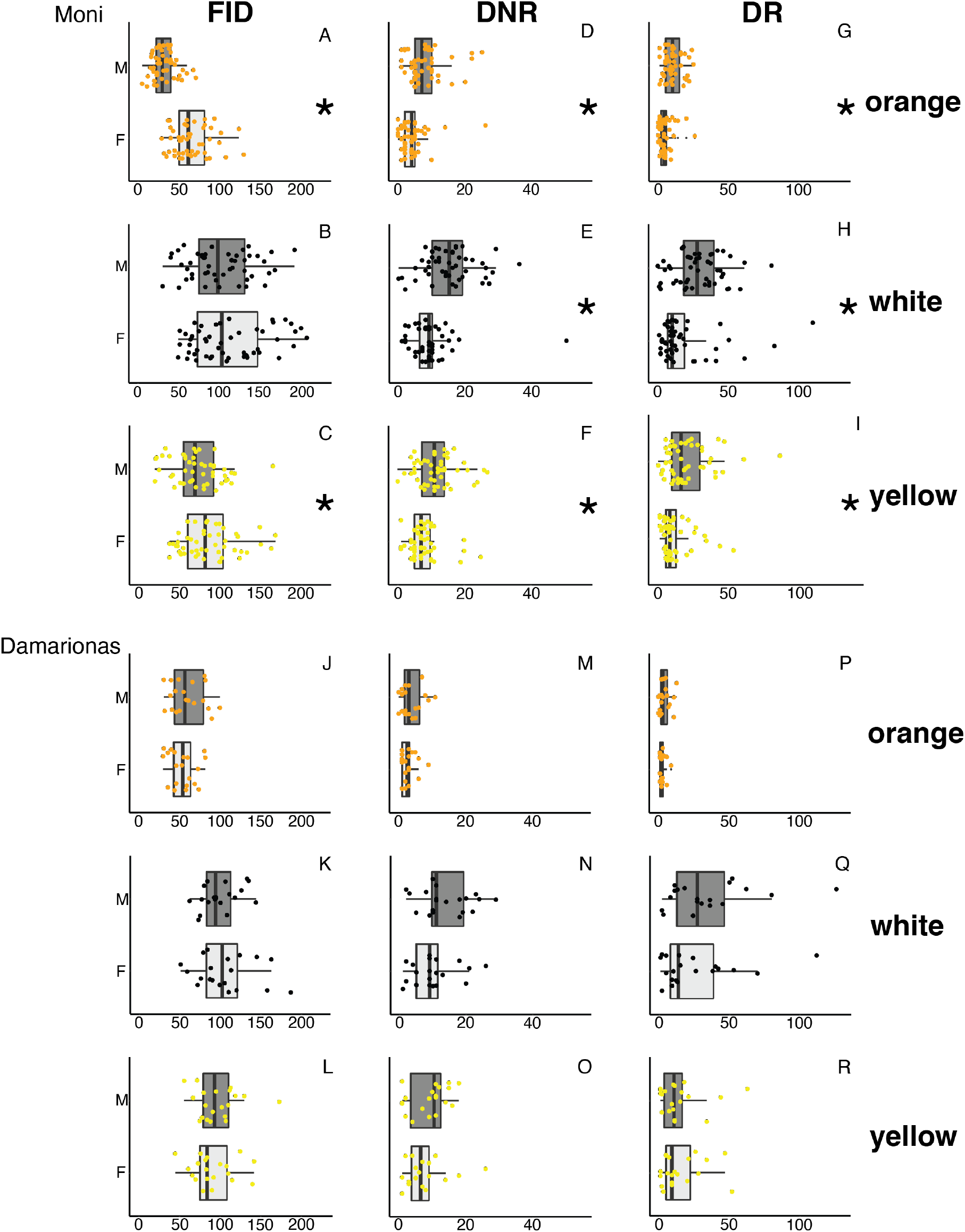
Color morph sex differences in flight initiation distance FID (A-C), distance to nearest refuge DNR (D-F, M-O), and distance to chosen refuge DR (G-I, P-R) at Moni and Damarionas. Asterisks indicate mean differences (alpha < 0.05).

At Damarionas, we did not observe any morph sex differences in any of the escape behaviors we measured (Figure 2 J-R). FIDs of the sexes were similar for the orange (t = −1.03, df = 38, p = 0.31), white (t = 0.68, df = 38, p = 0.49), and yellow morphs yellow (t = −0.7, df = 38, p = 0.49). DNRs did not significantly differ between the sexes for orange (t = −1.17, df = 38, p = 0.25), white (t = −1.61, df = 38, p = 0.14), or yellow morphs (t = −0.74, df = 38, p = 0.47). Finally, we detected no sex differences distance to the chosen refuge in orange (t = −1.42, df = 38, p = 0.17), white (t = −1, df = 38, p = 0.32), or yellow morphs (t = 0.18, df = 38, p = 0.85).

### Morph refuge selection

At Moni and Damarionas, orange morphs chose refuge in vegetation more often than the other morphs (Moni: *χ*^2^ = 19.12, df = 2, p < 0.001; Damarionas: *χ*^2^ = 13.5, df = 2, p = 0.001; Figure 3). At Moni, 23 orange morphs, 3 white morphs, and 8 yellow morphs chose refuge in vegetation, and 77 orange morphs, 97 white morphs, and 92 yellow morphs chose refuge in dry stone wall crevices. At Damarionas 10 orange morphs, 1 white morph, and 1 yellow morph chose refuge in vegetation, and 30 orange morphs, 39 white morphs, and 39 yellow morphs chose refuge in wall crevices. In general, all morphs chose refuge in the crevices of dry stone walls more often than vegetation (Figure 3).

**Figure 3.**
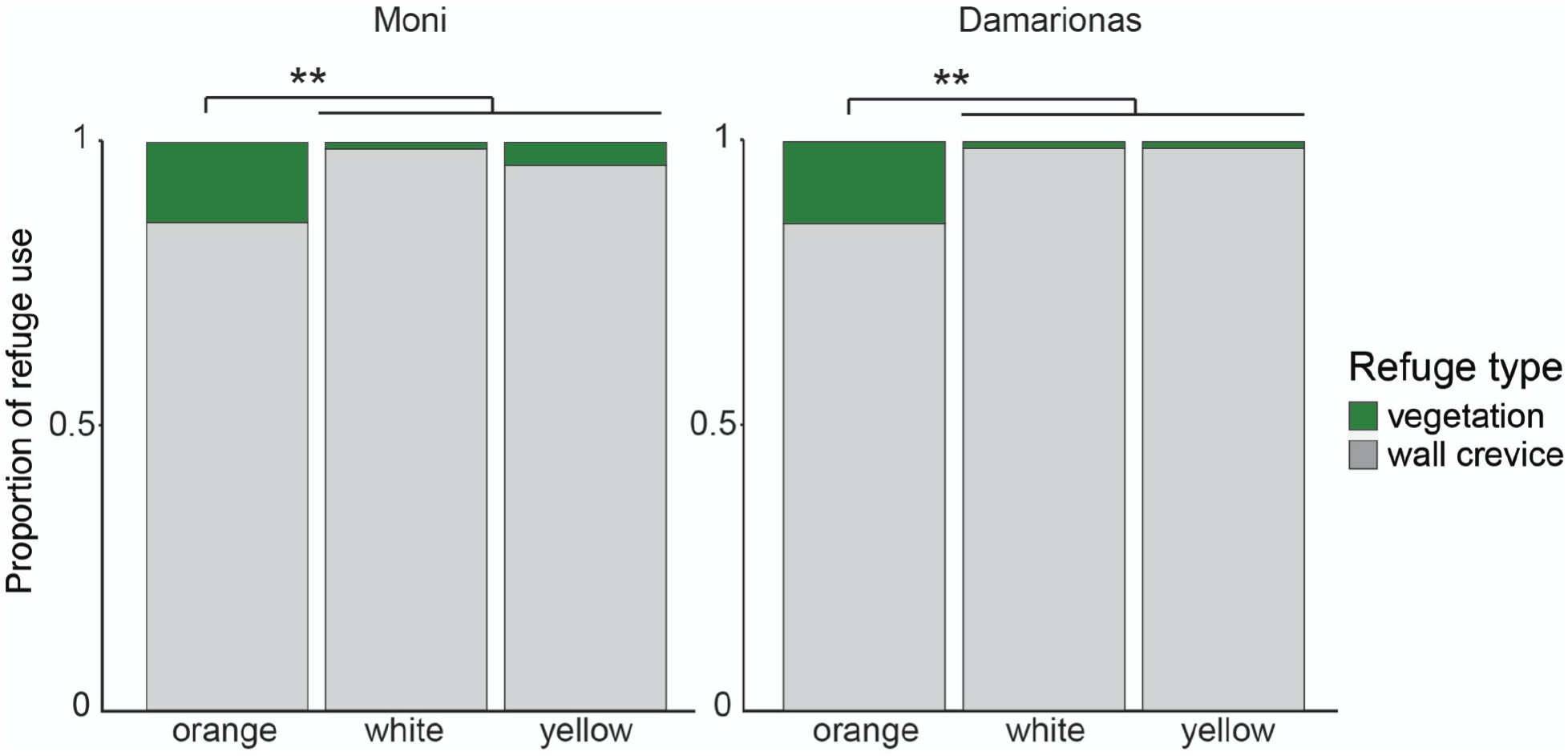
Proportion of chosen refugia by color morph at each field site. At both Moni and Damarionas, orange morphs used vegetation as a refuge more often than white and yellow morphs.

### Morph re-emergence behavior

We found consistent, across-site differences in morph re-emergence behavior. At Moni, color morphs differed significantly in the number of times they re-emerged from refuge within a 30-second period (Kruskal-Wallis chi-squared = 150.1, df = 2, p < 0.001). A post hoc Mann-Whitney U test revealed that all color morphs differed significantly from each other at Moni (Mann-Whitney U with Bonferroni correction, orange-white p < 0.001; white-yellow p < 0.001; yellow-orange p < 0.001). Color morphs also differed significantly in the number of times they re-emerged from refuge at Damarionas (Kruskal-Wallis chi-squared = 61.027, df = 2, p < 0.001). Similar to Moni, color morphs from Damarionas all differed significantly from each other in how often they re-emerged from hiding in refugia (Mann-Whitney U with Bonferroni correction, orange-white p < 0.001, white-yellow p < 0.001, yellow-orange p < 0.001).

At both Moni and Damarionas, orange morphs re-emerged from refuge the fewest times (Table 2). At Moni 6% of orange morphs re-emerged from refuge and only 10% re-emerged at Damarionas. When orange morphs did re-emerge from hiding, they took about twice as long to do so (on-average) compared to other morphs (Table 2). Yellow morphs re-emerged from hiding 36% of the time at Moni, and 53% of the time at Damarionas. Yellow morphs were quicker to re-emerge than orange morphs and took a similar amount of time to re-emerge as white morphs at both sites. Finally, white morphs re-emerged from hiding the most often compared to other morphs at both sites, with a re-emergence rate of over 90% at both sites (Table 2).

**Table 2.**
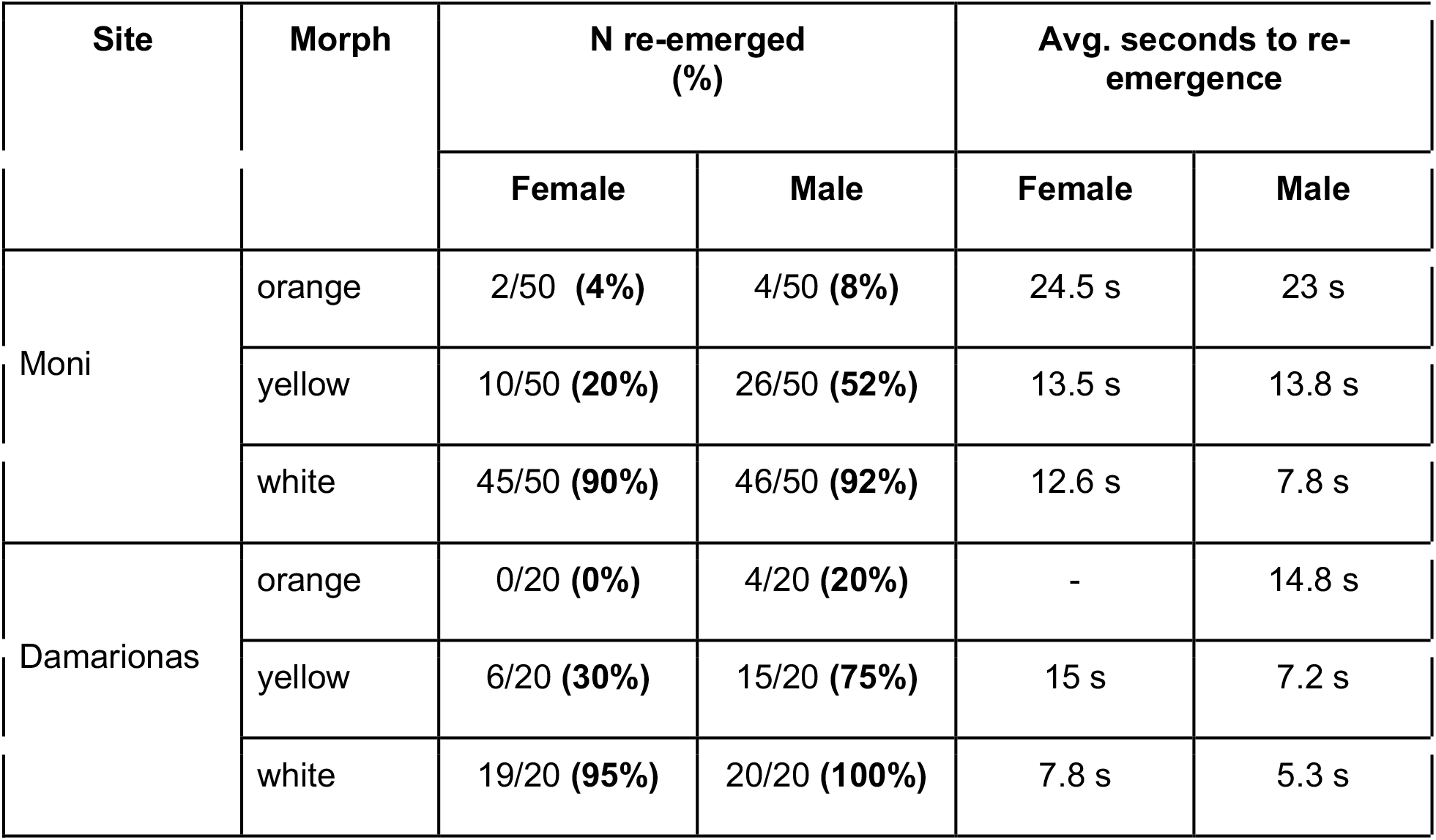
Morph re-emergence frequencies and time to re-emergence from both field sites. At both Moni and Damarionas, significantly fewer orange morphs re-emerged in a 30 second period after taking refuge than white or yellow morphs. Of individuals that did re-emerge, orange morphs took significantly longer to do so.

| Site | Morph | N re-emerged (%) |  | Avg. seconds to re-emergence |  |
| --- | --- | --- | --- | --- | --- |
|  |  | Female | Male | Female | Male |
| Moni | orange | 2/50 (4%) | 4/50 (8%) | 24.5 s | 23 s |
|  | yellow | 10/50 (20%) | 26/50 (52%) | 13.5 s | 13.8 s |
|  | white | 45/50 (90%) | 46/50 (92%) | 12.6 s | 7.8 s |
| Damarionas | orange | 0/20 (0%) | 4/20 (20%) | - | 14.8 s |
|  | yellow | 6/20 (30%) | 15/20 (75%) | 15 s | 7.2 s |
|  | white | 19/20 (95%) | 20/20 (100%) | 7.8 s | 5.3 s |

## DISCUSSION

Distinct color morph-correlated behavior may play a role in the evolution and maintenance of polymorphisms (Sinervo & Lively 1996; Punzalan et al. 2005; Gray & McKinnon 2007). In this study, we examined color morph differences in escape behavior to determine if color morphs have distinct strategies for avoiding predation. We found that co-occurring color morphs exhibit differences in their flight initiation distance, distance from nearest refuge, distance to chosen refuge, type of refuge they select, and re-emergence behaviors. We also found that color morph-correlated anti-predator behaviors were consistent in the two populations we sampled. Our results add to a growing body of evidence that color morphs in this species have evolved alternative life history traits and behavioral strategies (Brock et al. 2020; Brock et al. 2021a in review), which may play a role in color morph evolution and coexistence.

Visual predators may impose correlational selection on prey color and behavior (Forsman & Appleqvist, 1998), and influence color morph evolution, frequencies, and persistence in populations (Shigemiya 2004; Punzalan et al. 2005). Common wall lizards (*Podarcis muralis*) are a close relative to *P. erhardii* and have the same three monochromatic color morphs (Andrade et al. 2019). In *P. muralis*, orange morphs are more conspicuous to conspecifics and predators than yellow or white morphs (Pérez i de Lanuza & Font, 2015). In *P. erhardii*, orange, yellow, and white morphs have distinct and discernable spectra (Brock et al. 2020) and may also be discernible by predators with advanced visual systems such as birds. Birds are known to impose apostatic selection, or disproportionately consume common color morphs in a population (Allen 1972; Allen et al. 1998). This apostatic selection is a form of negative frequency-dependent selection that could promote polymorphism by favoring a rare morph, allowing it to persist in the population. Long-term monitoring of polymorphic populations is needed to discern whether color morph frequencies change following a frequency-dependent pattern in this species. It is also worth noting that *P. erhardii* male color morphs have somewhat distinct chemical signal profiles in their femoral pore exudate (Brock et al. 2020). Although not evaluated in this study, chemosensory cues may make certain color morphs more susceptible to detection by certain types of predators, particularly snakes and mammals. It is known that different types of predators have different effects on FID and tail-shedding behavior in *P. erhardii* (Brock et al. 2015), but how those differences pertain to color morph evolution and persistence is unknown. Further research on the visual and chemical ecology of Aegean wall lizards and their predators is needed to determine if certain color morphs are more conspicuous to predators, and under what circumstances.

Morph-correlated behaviors can be indicators of broader life-history differences and alternative strategies in color polymorphic species (Yewers et al. 2016). In color polymorphic *Podarcis* lizards, orange morphs tend to be restricted to vegetated habitats close to water (Pérez i de Lanuza & Carretero, 2018; Brock et al. 2021b in review). In our study, we found that orange morphs used vegetation as refuge more often, stayed the closest to refuge, and had shorter FIDs compared to other morphs. Orange morphs in *Podarcis* lizards may stay close to vegetation and refuge due to greater detectability by predators (Pérez i de Lanuza & Font, 2015), and may have shorter FIDs due to maintaining close proximity to refuge. Alternatively, orange morphs may develop associations with vegetated habitat and shorter FIDs in response to more frequent predator harassment. If orange morphs are more easily detectable by predators, then they may experience being chased more often and develop higher risk thresholds for fleeing, which would translate to shorter FIDs. Time spent fleeing cannot be spent on other necessary and fitness-enhancing activities such as feeding, thermoregulating, socializing, and mating. Orange morphs may remain close to vegetation and refuge to avoid detection and reduce the amount of time being chased, resulting in shorter FIDs.

Our results add support to the idea that wall lizard color morphs exhibit alternative behavioral strategies (Brock et al. 2021a in review). In this study, white and yellow morphs were observed further away from refuge and chose more distant refugia than orange morphs, which are facets of boldness. Boldness is often correlated with high levels of aggression toward conspecifics (Coleman & Wilson 1998; Reaney & Backwell 2007). White and yellow male morphs exhibited higher levels of bold and aggressive behaviors than orange male morphs in conspecific contest experiments (Brock et al. 2021a in review). Higher levels of boldness and aggression and larger body size are usually associated with dominance (Francis 1988; Sinervo & Lively 1996), and dominance can determine relative resource access and fitness among morphs. In this species, white and yellow morphs have higher levels of boldness and aggression (Brock et al. 2021 a), and the orange morph has larger head and body sizes (Brock et al. 2020). Morph-correlated behaviors and morphological traits likely offer context-specific advantages, resulting in somewhat equal population-level morph fitness over time and balancing the polymorphism. Our study is an important step in characterizing the color polymorphism in this species, and provides another piece to the puzzle of polymorphism evolution and maintenance. More research that focuses on morph relative fitness and trait values is needed to provide insight into the mechanisms regulating polymorphisms.

In the well-studied color polymorphic *Podarcis* species, the orange morph tends to be more behaviorally and phenotypically distinct from the yellow and white morphs (Huyghe et al. 2007; Huyghe et al. 2009; Abalos et al. 2016; Brock et al. 2021a in review), and the yellow and white morphs tend to be more similar to each other. Morph-assortative mating could minimize the break-up of coadapted gene complexes and generate linkage disequilibrium built up by correlational selection (McKinnon & Pierotti 2010), resulting in phenotypic divergence between morphs. Homomorphic pairs tend to be more common than heteromorphic pairs in several *Podarcis* species (Huyghe et al. 2010; Pérez i de Lanuza et al. 2013; K.M. Brock unpublished data). Further, white and yellow morphs tend to be found in more similar habitat and behave more similarly to each other than the orange morph (Huyghe et al. 2007; Brock et al. 2021a, 2021b in review). Thus, it is possible that white-yellow is a more common heteromorphic mate pairing than orange-white or yellow-orange pairings. If there is non-random mating in populations of color morphs, it could lead to deeper genetic divergences between certain morphs (Huyghe et al. 2010), with potential effects on phenotypic divergence. More work on color morph genomes is needed to understand the genetic basis of behavioral and phenotypic differences among color morphs and their implications in morph evolution and maintenance.

We demonstrated that color morph-correlated behaviors are repeatable across distinct populations, a detail that is often assumed in other behavioral studies concerning color polymorphic species. Although color morph differences in behaviors such as boldness (Yewers et al. 2016), aggression (Abalos et al. 2016; Yewers et al. 2016), and mating strategies (Sinervo & Lively 1996), are often observed in color polymorphic species, these differences are rarely demonstrated to be true for more than one population. Confirming consistency in alternative behavioral phenotypes is essential when trying to understand drivers of phenotypic variation.

Future work should also explore color morph behavioral differences in populations with different ecological contexts. Whether or not different predation regimes have different effects on color morph behaviors, life histories, or persistence in populations remains an open question in this and many other color polymorphic species.

## ACKNOWLEDGEMENTS

We would like to thank Dr. Panayiotis Pafilis at the National and Kapodistrian University of Athens for his assistance with research permits.

